# Getting over ANOVA: Estimation graphics for multi-group comparisons

**DOI:** 10.64898/2026.01.26.701654

**Authors:** Zinan Lu, Jonathan Anns, Yishan Mai, Rou Zhang, Kahseng Lian, Nicole MynYi Lee, Shan Hashir, Lucas Wang Zhuoyu, Yixuan Li, A. Rosa Castillo Gonzalez, Joses Ho, Hyungwon Choi, Sangyu Xu, Adam Claridge-Chang

**Affiliations:** Program in Neuroscience and Behavioural Disorders, Duke-NUS Medical School, Singapore; Department of Statistics and Data Science, National University of Singapore, Singapore; Institute for Molecular and Cell Biology (IMCB), Agency for Science, Technology and Research (A*STAR), Singapore, Singapore; Department of Medicine, Yong Loo Lin School of Medicine, National University of Singapore, Singapore; Singapore Lipidomics Incubator, Life Sciences Institute, National University of Singapore; Department of Physiology, National University of Singapore, Singapore

## Abstract

Data analysis in experimental science mainly relies on null-hypothesis significance testing, despite its well-known limitations. A powerful alternative is estimation statistics, which focuses on effect-size quantification. However, current estimation tools struggle with the complex, multi-group comparisons common in biological research. Here we introduce DABEST 2.0, an estimation framework for complex experimental designs, including shared-control, repeated-measures, two-way factorial experiments, and meta-analysis of replicates.

## Introduction

For decades, statisticians have pointed out the limitations of *null-hypothesis significance testing* (NHST) as a method for data analysis. The NHST framework turns research decision-making into a misleading dichotomy between so-called ‘significance’ and ‘non-significance’ (Berkson, 1942; Cohen, 1994; Nuzzo, 2014; Wasserstein, R. L., Schirm, A. L. & Lazar, N. A. eds., 2019). As a result, researchers often place undue confidence in NHST test results (Button et al., 2013; Halsey et al., 2015), contributing to poor experimental reproducibility (Collaboration, 2015; Errington et al., 2021; Prinz et al., 2011). The emphasis on this dichotomy also discourages researchers from engaging in the critical task of effect quantification (Cumming & Calin-Jageman, 2017) (**Supplementary Note 1**).

An alternative to NHST is estimation statistics, a data analysis framework that comprises effect size estimation, precision intervals, data graphics and meta-analysis (Altman et al., 2000; Borenstein et al., 2009; R. J. Calin-Jageman & Cumming, 2019; Claridge-Chang & Assam, 2016; Singh Chawla, 2017). The estimation framework has the potential to almost entirely replace NHST. However, realizing this transition will require the implementation of estimation methods, accessible software and data graphics appropriate for all experimental types, including complex designs (**Supplementary Note 2**).

### Conventional multi-group analysis is unfocused

In addition to the limitations of NHST in simple comparisons, multi-group experiments present greater challenges. Conventional multi-group analysis uses omnibus testing via Analysis of Variance (ANOVA, (Fisher, 1925)) and its variants, followed by multiple comparisons (Dunn, 1961). However, ANOVA tests a null hypothesis that is inherently unfocused, assuming “everything is the same” (all group means are equal). A rejection of this proposition provides no specific information about which groups differ, in which direction, or by how much (Rosenthal & Rosnow, 1985). Consequently, researchers must embark on a second analytical step that requires multiple comparisons. Even a modest number of groups (*g*) requires the analyst to test *m* hypotheses; with *m* = *g*(*g* − 1)/2 pairwise comparisons, the number of tests sharply increases. For example, a 6-group experiment requires the testing of 15 hypotheses. Thus, in a typical multi-group experiment, what may have begun as a focused research question sprawls into a complex web of subsidiary tests. Multiplicity is usually managed with adjustments such as Bonferroni correction, but these undermine the statistical power and make it harder to detect true effects (García-Pérez, 2023) (**Supplementary Note 3**).

Estimation graphics, using the Cumming-plot design, encourage the researcher to limit the comparisons in multi-group analysis, for example, by comparing each test group only to a single control (Cumming & Calin-Jageman, 2017; Dunnett, 1955). For *g* groups, this means focusing on *g* − 1 effect sizes, *e*.*g*. instead of 15 *p* values, a 6-group experiment considers only 5 effect sizes. Multi-group estimation graphics show the effect magnitudes and precision estimates—the information most relevant for understanding biological systems. We also remark that existing effect-size software tends to only facilitate the analysis of simpler two-group experimental designs. Here we introduce DABEST 2.0, software that extends the applicability of estimation graphics to more complex multi-group experiments, including longitudinal studies, two-way factorial designs, multi-group binary data, and meta-analyses of internal replicates.

## Results

### Repeated-measures estimation graphics display effect trajectories

DABEST 1.0 was introduced with an estimation graphic for the multi-group data normally analysed via one-way ANOVA (Ho et al., 2019). With DABEST 2.0, we first extended this functionality to repeated-measures experiments. Throughout this article, we demonstrate estimation graphics with fictional scenarios. Consider a time-course study on the total sleep time (TST) of insomnia patients during drug treatment. The researcher records each patient’s sleep period at baseline and each day after treatment (**Figure 1a**). To analyse the data with a repeated-measures ANOVA, the researcher first calculates the *F* statistic and its *p-*value. If deemed ‘significant’, the data are analysed with several pairwise *t*-tests: each time point is compared against all others, generating several T-values (test statistic) and multiple-comparison-adjusted *p* values (**Figure 1b, Supplementary Table 2**).

**Figure 1.**
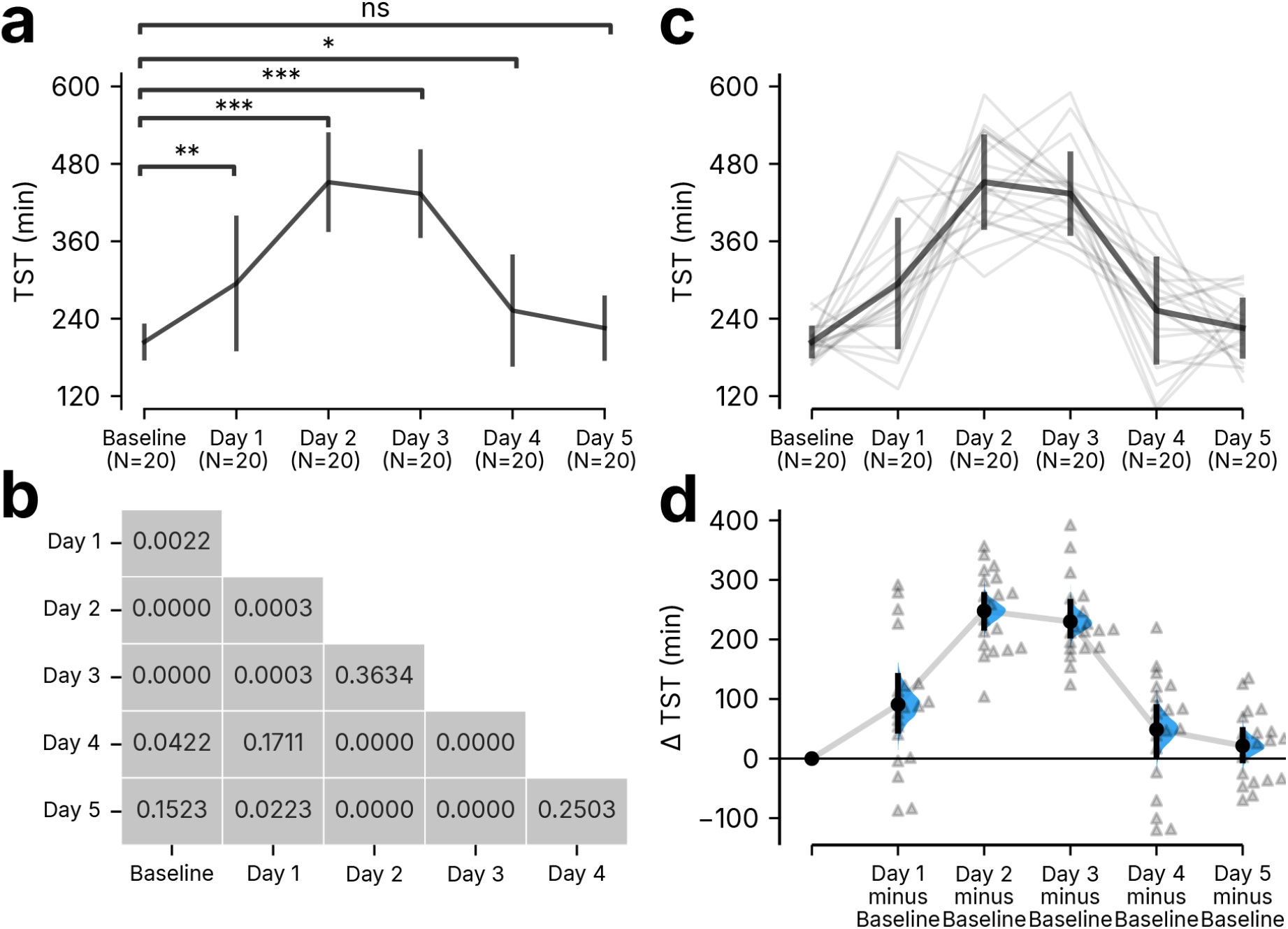
Estimation graphics for multiple group experimental designs. **a**, A conventional repeated measures plot of total sleep time (TST) in minutes (min) following treatment with an insomnia drug. One-way ANOVA with *post hoc* testing was used to generate corrected *p* values for all group comparisons. The *p* values for the comparison of each test day versus baseline are represented by asterisks (***: p<0.001; **: 0.001<p<0.01; *: 0.01<p<0.05, ns: p≥0.05). **b**, The full *p* value table for the 15 comparisons. **c–d**, Repeated-measures plot of the same data as in **a**. Panel **c** shows individual subjects’ data as connected lines: grey lines indicate the individual paired samples, while the black line indicates the mean and standard deviation.. Panel **d** shows the bootstrapped distributions of the differences between test days and baseline. Black vertical bars indicate 95% confidence intervals. Triangles indicate individual deltas.

The effect-size alternative is the repeated-measures estimation plot (**Figure 1c–d**). These two panels serve distinct purposes: the top panel (**Figure 1c**) displays observed values and their dispersion (also shown as standard-deviation bars), while the bottom panel displays effect sizes and their precision (**Figure 1d**), shown as error curves and confidence-interval bars, which will narrow with increasing sample size. In large-sample experiments, observed values may still appear dispersed while the effect curve simultaneously reveals narrow, precise estimates; presenting both descriptive and inferential data together conveys the complete picture of an analysis.

The effect size plot allows the researcher to focus on the relevant time point versus the baseline comparison. The effect sizes and their confidence intervals (CIs), calculated via bootstrapped resampling (DiCiccio & Efron, 1996; Efron, 1979) (**Supplementary Note 4**), are represented by the dots (effect size), vertical black lines (95CI), and the full distributions (half-violin curves). The estimation framework shifts the focus to the magnitude of the intervention effect size (compared to Figure 1a). Thus, the researcher can clearly see that total sleep time slightly increased on Day 1 (90.8 min [95CI 46.8, 139.0]), greatly increased by Day 2, and plateaued by Day 3 (247.8 min [95CI 219.5, 274.6], and 223.0 min [95CI 206.3, 263.1], respectively), before declining over the subsequent days.

### Two-factor designs: delta-delta analysis

In complex experiments with two independent variables, researchers conventionally rely on two-way ANOVA (Fisher, 1935), often including an interaction term for interdependent effects. This approach still requires *post-hoc* tests for explicit group-to-group comparisons. DABEST 2.0 simplifies this analysis by introducing the delta-delta effect size, which focuses on the net effect of one primary variable within the context of a secondary variable (**Supplementary Note 5**).

Consider an experiment designed to test a drug’s efficacy in treating test animals with a disease-causing mutation. This hypothetical experiment measures survival in both mutant (M) and wildtype (W) subjects given drug and placebo treatments. Key questions include (1) By how much does the drug improve survival in mutation carriers? (2) What placebo effect exists in mutation carriers? and (3) What is the drug’s net effect after accounting for placebo effects?

Two-way ANOVA (**Supplementary Table 3**) shows a significant interaction effect (*F*(1,196) = 26.56, *p* = 6.21e^−7^), indicating that treatment effects depend on subject genotype, and Tukey’s test is then performed (**Figure 2a**). In this case, of the six possible comparisons, four yield significant differences. Despite this, how much the drug specifically benefits mutation carriers remains unclear, making interpretation challenging.

**Figure 2.**
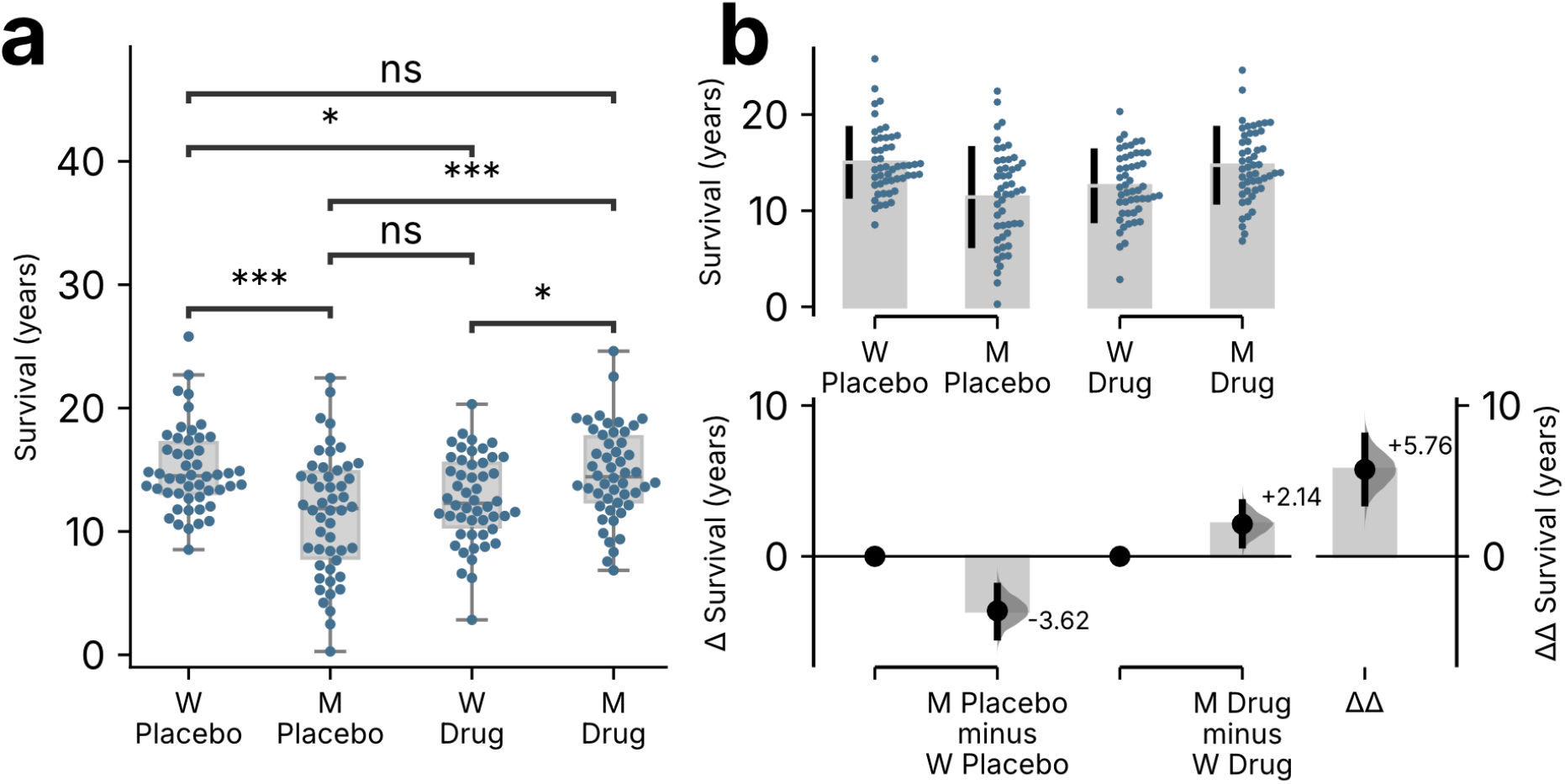
Estimation statistics for drug × genotype effects. **a**, A swarm and box plot of survival effects. The box shows the interquartile range (25th to 75th percentiles) with a horizontal line at the median (50th percentile). Vertical lines (whiskers) extend to 1.5 times the interquartile range from the upper and lower quartiles. Horizontal brackets with stars indicate the level of significance for comparisons with post-hoc Tukey’s test (***: p<0.001; **: 0.001<p<0.01; *:0.01<p<0.05). **b**, Estimation graphic of the survival effects between mutant (M) and wild type (W), calculated for the placebo and drug-treated groups; the delta-delta effect was calculated by subtracting the two primary deltas. The top panel is a swarm plot of the samples alongside a vertical bar, the gap indicates the mean of the sample, while the lines show the standard deviations. In the bottom panel, the left y-axis shows the bootstrapped distributions of the mean differences; the right y-axis shows the delta-delta effect size. Black vertical bars indicate 95% confidence intervals.

Delta-delta analysis clearly depicts the two-way drug × genotype effect and reveals three key points (**Figure 2b**). First, mutation carriers given the placebo show reduced survival, by approximately –3.62 years [95CI –5.29, –2.01]; second, carriers given the drug treatment show improved survival, by about 2.1 years [95CI 0.76, 3.61]; and third, according to the delta-delta calculation (obtained by subtracting the placebo delta from the drug delta; **Methods**), which indicates the drug’s net effect, accounting for all background effects. Where an ANOVA user would conclude only that a statistically significant interaction exists, a delta-delta user reads directly that, in carriers, the drug improved survival by approximately 5.76 years [95CI 3.60, 7.89] (**Supplementary Note 6**).

This approach should prove useful for researchers seeking to quantify specific treatment effects via 2×2 designs, using either original units or standardized effect sizes (**Methods**). For either paired data or independent groups, delta-delta graphics account for non-specific, background effects and distil a multifactorial problem into a single, focused effect size.

### Differences of proportions

The NHST framework commonly employs Fisher’s exact test or the Chi-squared test to analyse binary (or ‘proportional’) data (Kim, 2017). A systematic review of 70 published articles found that most Fisher’s exact test results were reported without any visualization of precision, and not one reported a numerical effect size (**Supplementary Table 4**).

To illustrate the differences between NHST and estimation analyses of proportional data, consider an experiment to test the efficacy of a drug in an animal model of spontaneous seizures. The drug is administered for 3 days, and seizure occurrence is scored as a binary outcome per animal (**Supplementary Table 5**). The researcher compares the seizure occurrence on day 3 in the placebo and drug-treated groups. Instead of a standard bar chart, which often omits error and effect statistics (**Figure 3a**), DABEST 2.0 provides a proportion plot incorporating error bars and the effect size. Compared to the placebo treatment, the drug reduced the number of spontaneous seizures in the model animals by 68% [95CI 53, 83] on day 3 (**Figure 3b**). DABEST 2.0 can also draw Sankey plots to illustrate changes in repeated-measures binary data (blogmaster_sev, 2021; Kosara, 2010). For example, changes in seizures over the 3 days relative to baseline can be visualized for the placebo (+5% [95CI –10, +18]) and drug groups (−60% [95CI −45, −78]) (**Figure 3c**). For reporting standardized effect sizes in binary data, DABEST can calculate Cohen’s *h* (**Methods**).

**Figure 3.**
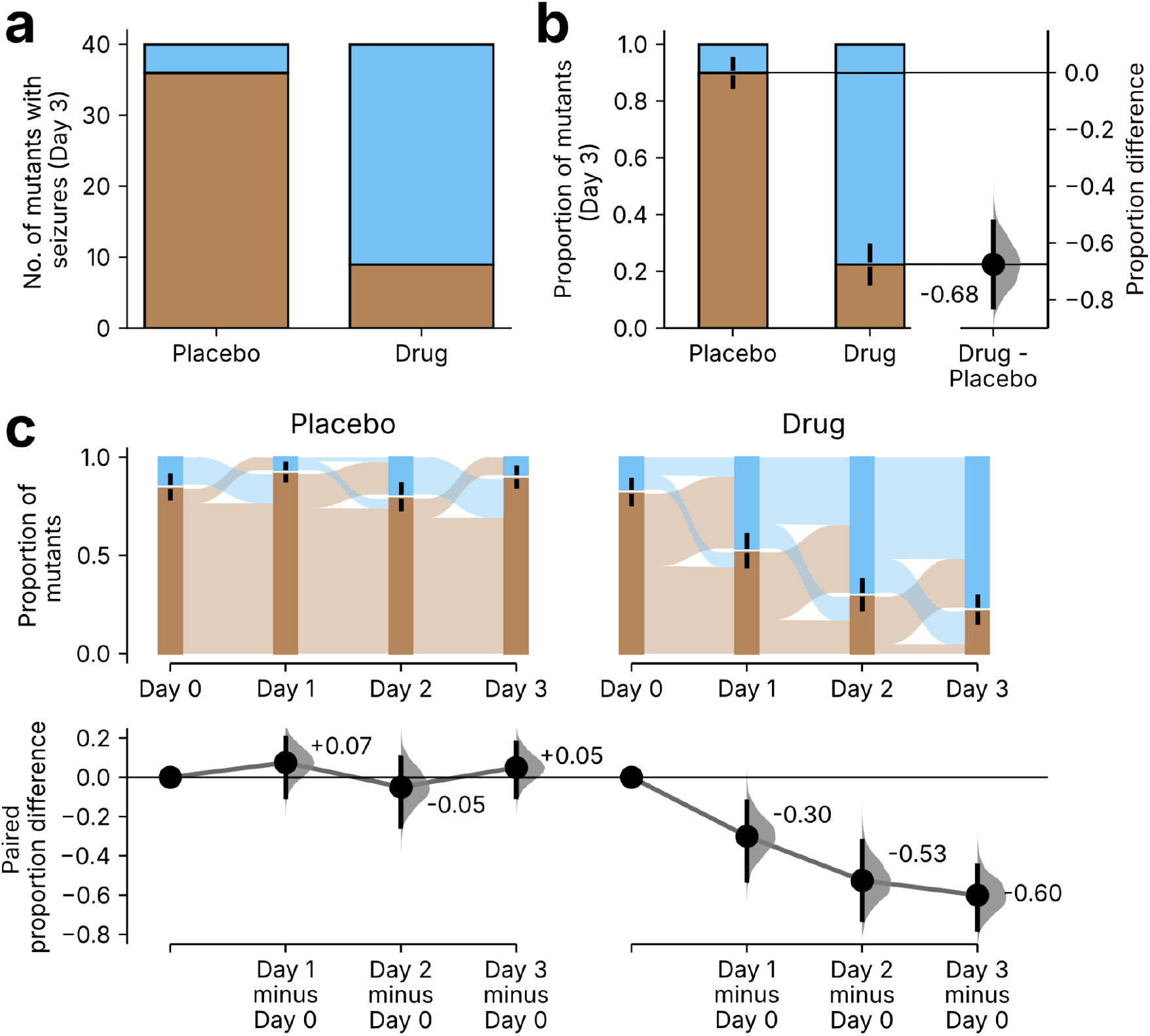
Graphical analyses of proportions and their differences. **a**, A bar chart that visualizes proportions without error bars or an effect size. **b**, Proportion plot generated using DABEST 2.0. The brown section of each bar represents the proportion of animals with seizures, while the blue section represents animals without seizures. **c**, Sankey-style estimation plots for repeated measures of proportions. The brown and blue bars illustrate the proportion of animals with and without seizures, respectively. The strip between each Sankey bar denotes categorical changes. Mean and ± standard deviation are represented by black vertical bars. Effect-size distributions are plotted as half-violin curves on either the right y-axis (**b**) or the x-axis (**c**, lower panel), alongside each mean (black dot) and 95% confidence interval (black line).

### Simple meta-analysis of replicated experiments

Meta-analyses calculate more precise effect estimates, and resolve contradictions between studies (Glass, 1976; Gurevitch et al., 2018; O’Rourke, 2007). However, researchers rarely meta-analyse data from internally replicated experiments.

Consider evaluating a drug’s effect on total sleep time in insomnia patients in three pilot trials (**Figure 4**). Two replicates show substantial effect sizes, while one does not. Without a way to synthesize this, researchers must either report only one experiment or combine all data into a single mega-analysis. Selective reporting conceals experiments, while clumping data lacks transparency about the experimental progression (including possible inter-experiment modifications), magnifies errors when baselines vary, and can encourage *p*-hacking (Head et al., 2015).

**Figure 4.**
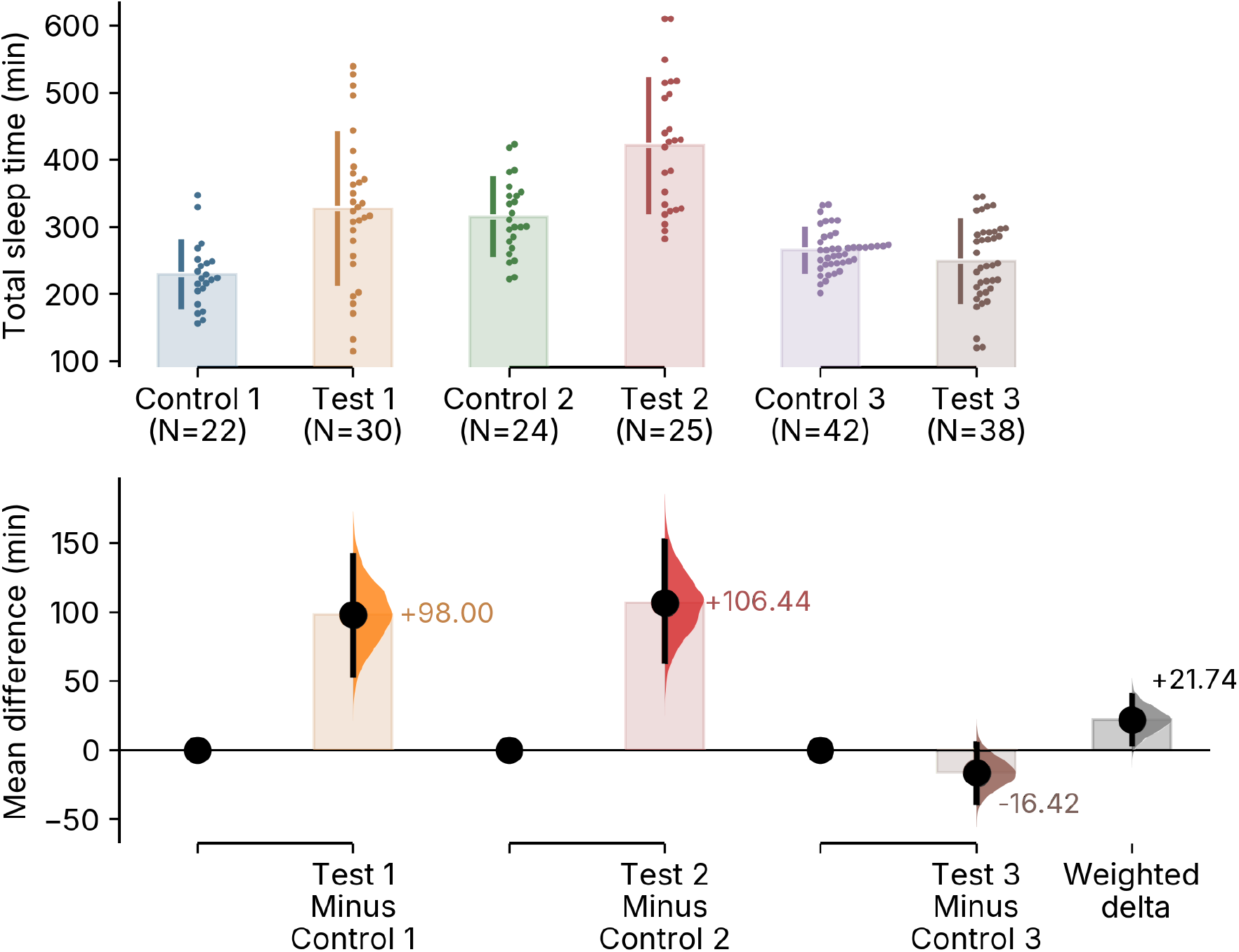
Example mini-meta-analysis of data from 3 two-group experiments. **Top panel**. Swarm plots of observed values from three experiments, with a vertical bar indicating the mean and standard deviation. **Lower panel**. The weighted-average delta resolves differences between replicates with positive effect sizes (Test 1 minus Control 1, +98.00 [95CI +54.25, +140.34]; Test 2 minus Control 2, +106.44 [95CI +64.44, +151.23]) and a negative effect size (Test 3 minus Control 3, –16.42 [95CI –37.69, 4.14]). The mean differences of the 3 experiments: half-violin curves show the bootstrapped distribution of differences, and vertical bars show the 95% confidence intervals. The right-most column shows the weighted-average mean difference (Δ_w_ = +21.74 [95CI +4.51, +39.32]).

To address these issues, several groups have argued for the use of a small-scale meta-analysis or ‘mini-meta’ (Braver et al., 2014; Cumming, 2014; Goh et al., 2016; Maner, 2014) (**Supplementary Note 7**). To date, visualizing a series of replicated experiments still requires the use of standard meta-analysis tools (Goh et al., 2016). DABEST 2.0 introduces easy mini-meta-analysis, where users can display multiple instances of a series of experiments, their individual effect sizes, and the summary effect (**Figure 4**). The summary effect is calculated as a weighted delta using pooled replicate variances and a fixed-effects model suitable for replicates of similar experiments (**Methods**). Visualizing individual replicates shows both intra- and inter-experimental variations, while the averaged mean difference provides the overall most plausible effect size. The adoption of mini-meta promotes the disclosure of seemingly contradictory results, encourages transparent reporting, enhances reproducibility, and improves estimate precision.

## Conclusion

DABEST 2.0 expands the applicability of estimation graphics with new tools for analysing repeated-measures experiments, two-way designs, and binary/proportional data. As well as providing alternatives to three major classes of significance tests, DABEST 2.0 also provides a solution for handling replicated experiments.

DABEST 2.0 is available as Python and R packages (with comprehensive documentation and tutorials) and as a web application, https://estimationstats.com. By providing robust estimation solutions for common multi-group experiments, DABEST 2.0 facilitates the transition from significance testing to more meaningful quantitative analysis.

## Supplementary Notes

1. A common concern is that effect-size interpretation is subjective. We consider this a strength rather than a limitation (Wasserstein et al., 2019): whether an effect is meaningful should ultimately be resolved by scientific judgment within each field (Cohen, 1994; Gelman & Hennig, 2017). A field may develop its own guidelines for judging effect sizes; where such norms are lacking, Cohen’s conventional descriptors (small, medium, large) serve as a provisional guide (Cohen, 1988). Estimation encourages scientists to develop context-specific understanding of effect sizes in their systems; such expert judgment is essential to data analysis and interpretation (Wasserstein, R. L., Schirm, A. L. & Lazar, N. A. eds., 2019).
2. Until recently, ready-to-use software tools for estimation graphics were not widely available; this changed with the release of the DABEST suite, providing tools suitable for both scripting and point-click workflows: [DABEST](https://acclab.github.io/DABEST-python/) for Python, [dabestr](https://cran.r-project.org/web/packages/dabestr/index.html) for R, and estimationstats.com (R. Calin-Jageman & Cumming, 2024; Ho et al., 2019). This advance was reinforced by changes in journal policy (Bernard, 2019, 2021; Elkins et al., 2022) and a growing number of freely available software packages, including ESCI for R and Jamovi (R. Calin-Jageman & Cumming, 2024), *t*-tests with effect sizes in JASP (JASP Team, 2024), and [Durga](https://academic.oup.com/jeb/advance-article/doi/10.1093/jeb/voae073/7689073) for R (Khan & McLean, 2024) (**Supplementary Table 1**).
3. Strong adjustments such as Bonferroni correction undermine statistical power, increasing false negatives at the cost of reducing false positives (García-Pérez, 2023; Greenland, 2021; Hooper, 2025; Rothman, 1990). The individual confidence intervals reported by DABEST are computed marginally for each pairwise comparison, without correction for multiple testing. Users concerned about joint inference can widen their confidence interval beyond 95% (Benjamin et al., 2017).
4. Bootstrap resampling is used due to its computational simplicity, robustness for small samples and skewed/non-normal samples, and for allowing for calculation of the effect size and error-distribution curve without assumption of the underlying sample distribution, thus allowing for accurate estimation (Hesterberg, 2011; Horowitz, 2019). For confidence intervals, DABEST calculates bias-corrected and accelerated (BCa) bootstrap, which provides more accurate coverage than standard percentile intervals, particularly for small or skewed samples.
5. The delta-delta effect size estimates the same quantity as a two-way ANOVA interaction (the difference between two differences) but whereas ANOVA reduces this to an F-statistic and p-value, delta-delta displays it as an effect magnitude with a confidence interval. Where ANOVA’s interaction test asks only “is there any interaction?”, delta-delta estimates how much more or less an intervention works within each group context, reframing a multifactorial question into a directly interpretable, scientifically meaningful magnitude. In the general spirit of contrast analysis, a delta-delta plot allows the researcher to ask focused questions on the main driver variable as well as the differentials by the secondary variable in a quantitative manner.
6. Where an ANOVA user would have concluded that there is a statistically significant interaction between genotype and treatment (*p* = 6.21e-7) requiring further post-hoc testing, a delta-delta user concludes that the drug improved survival by 5.76 years [95% CI: 3.60, 7.89] in mutation carriers—an effect size that substantially exceeds the 2-year threshold typically considered clinically meaningful, with the confidence interval indicating that even the most conservative estimate represents substantial clinical benefit. Effect size estimation provides more information than NHST while maintaining statistical rigour.
7. A lack of internal replication is thought to contribute to the reproducibility crisis, with over 80% of researchers citing insufficient within-lab replication as a factor in irreproducibility (Baker, 2016). Compounding this issue is the fact that ‘negative’ replicates often remain unpublished—a well-known phenomenon known as ‘the file-drawer problem’ (Simonsohn et al., 2014).

**Supplemental Table 1.**
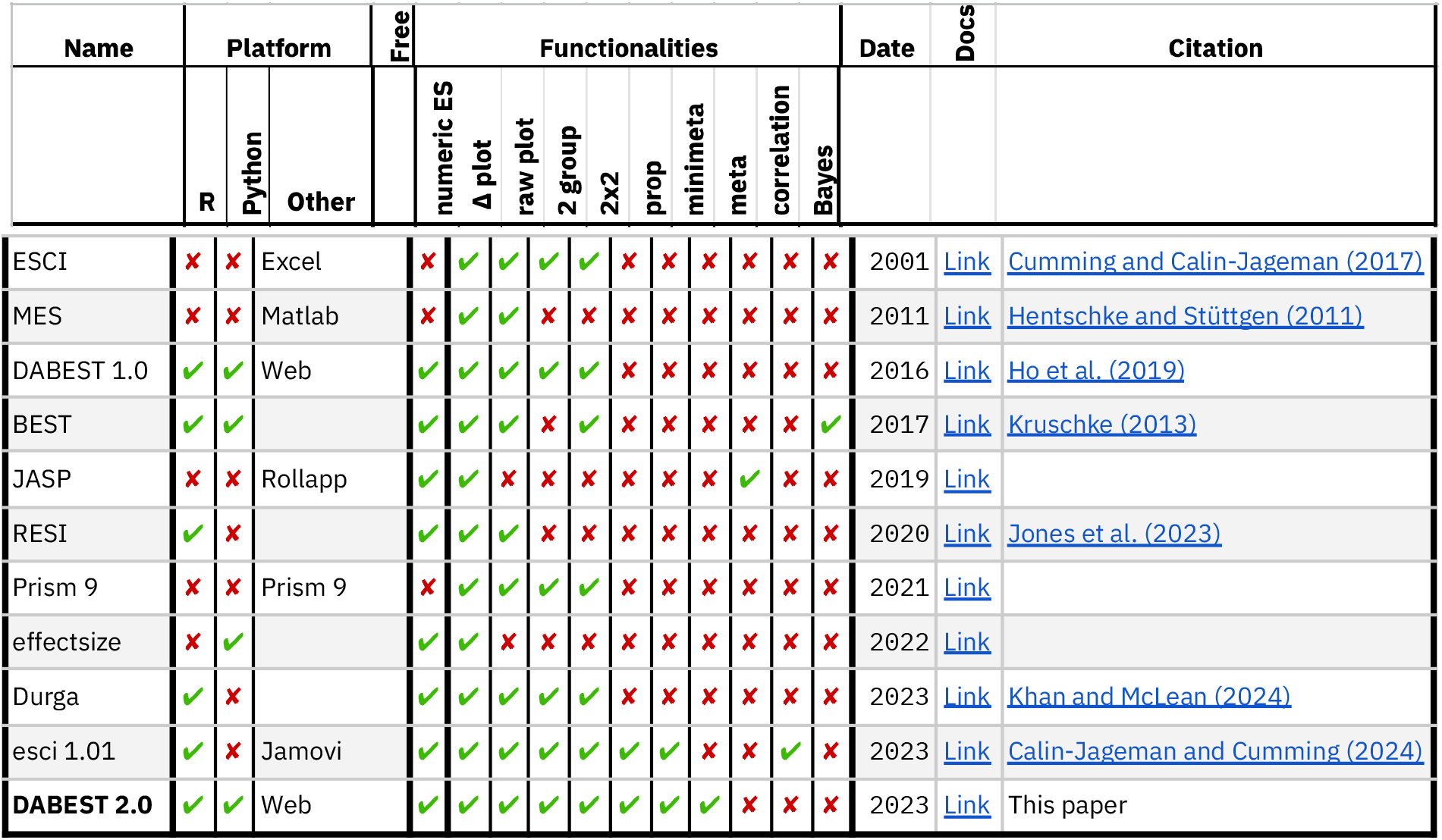
Comparison between major packages with support for effect size calculation and visualization. Comparison of major packages (ESCI (Cumming & Calin-Jageman, 2017), MES (Hentschke & Stüttgen, 2011), DABEST 1.0 (Ho et al., 2019), BEST (Kruschke, 2013), JASP, RESI (Jones et al., 2023), Prism 9, effectsize, Durga (Khan & McLean, 2024), esci 1.01 (R. Calin-Jageman & Cumming, 2024) and DABEST 2.0), across platform compatibility (R, Python, Excel, Matlab, Jamovi, web-based), cost (free vs. commercial), statistical capabilities (effect size metrics, group comparisons, proportions, meta-analysis, Bayesian methods), and visualization features (raw data plots, estimation plots). Date indicates year of release or major version. DABEST 2.0 is highlighted as the current package with comprehensive cross-platform support

**Supplemental Table 2.** Statistical analysis of the data presented in Figure 1A-B. Here, multiple pairwise t-tests are performed with Benjamini/Hochberg FDR correction.

| Group 1 | Group 2 | T statistic | Adjusted p-value | Reject H<sub>0</sub> |
| --- | --- | --- | --- | --- |
| Baseline | Day 1 | -3.7655 | 0.0022 | TRUE |
|  | Day 2 | -16.6801 | 0.0000 | TRUE |
|  | Day 3 | -15.7577 | 0.0000 | TRUE |
|  | Day 4 | -2.3300 | 0.0422 | TRUE |
|  | Day 5 | -1.6192 | 0.1523 | FALSE |
| Day 1 | Day 2 | -4.7033 | 0.0003 | TRUE |
|  | Day 3 | -4.7597 | 0.0003 | TRUE |
|  | Day 4 | 1.5068 | 0.1711 | FALSE |
|  | Day 5 | 2.6785 | 0.0223 | TRUE |
| Day 2 | Day 3 | 0.9312 | 0.3634 | FALSE |
|  | Day 4 | 6.9478 | 0.0000 | TRUE |
|  | Day 5 | 10.9465 | 0.0000 | TRUE |
| Day 3 | Day 4 | 8.6813 | 0.0000 | TRUE |
|  | Day 5 | 11.6226 | 0.0000 | TRUE |
| Day 4 | Day 5 | 1.2302 | 0.2503 | FALSE |

**Supplemental Table 3.** An ANOVA table for the 2×2 design drug experiment. There is a significant interaction effect between Genotype and Treatment, indicating that the treatment effect is specific to a genotype. Tukey’s post-hoc HSD test was therefore performed to determine the specific locations of the significant effects, shown in Figure 2a.

|  | df | sum_sq | mean_sq | F | PR(>F) |
| --- | --- | --- | --- | --- | --- |
| **C(Genotype)** | 1 | 27.33 | 27.33 | 1.75 | 0.19 |
| **C(Treatment)** | 1 | 9.48 | 9.48 | 0.61 | 0.44 |
| **C(Genotype) x C(Treatment)** | 1 | 414.31 | 414.31 | 26.56 | 6.21E-07 |
| **Residual** | 196 | 3057.44 | 15.6 |  |  |

**Supplemental Table 4.** Types of visualizations and tests used to showcase proportion related data in the literature. The first search (yielding 40 articles) used the keywords “Fisher’s exact test” in PubMed and Google Scholar from 2000 to 2024. These articles were further categorized based on the representation of how the Fisher’s test results were conveyed, with or without visualizations. The second search (yielding 30 articles) focused on the keyword “Proportion” in PubMed and categorized the presence or absence of error bars/confidence intervals as well as the types of statistical tests that were performed. Details of the literature review can be found in the Methods.

| Search type | Type of test | Visualization | Error | Count | % | Effect sizes |
| --- | --- | --- | --- | --- | --- | --- |
| **"Fisher's Exact test"** | Fisher's | Graphics | N | 19 | 47.5 | 0 |
|  |  |  | Y | 10 | 25.0 | 0 |
|  | Fisher's | No Graphics | N | 6 | 15.0 | 0 |
|  |  |  | Y | 5 | 12.5 | 0 |
| **"Proportion"** | Fisher's | Graphics | N | 5 | 16.1 | 0 |
|  |  |  | Y | 3 | 9.7 | 0 |
|  | Fisher's | No Graphics | N | 1 | 3.2 | 0 |
|  |  |  | Y | 4 | 12.9 | 0 |
|  | Mixed | Others | N | 12 | 38.7 | 0 |
|  |  |  | Y | 6 | 19.4 | 0 |

**Supplemental Table 5.** Example seizure occurrence drug trial data. Number of animals with spontaneous seizures across 3 days of placebo/drug administration. A binary seizure/no seizure scoring system was adopted, thus yielding binary data. Day 0 refers to the baseline prior to placebo/drug administration. n=40 for each group.

|  | Day 0 | Day 1 | Day 2 | Day 3 |
| --- | --- | --- | --- | --- |
| **Placebo** | 34 | 37 | 32 | 36 |
| **Drug** | 33 | 21 | 12 | 9 |

## Methods

### Data simulation

Data analysed in this paper were simulated using the SciPy, NumPy, and Pandas packages. All code necessary to reproduce the results with DABEST-python is available in a separate [Jupyter Notebook](https://drive.google.com/file/d/1J50LQpyFMg3Cm-DLQogCaL-VmVSuHNem/view?usp=drivesdk).

### Effect size calculations

#### Bootstrap resampling and confidence interval estimation

In all examples in this paper, 5000 pairs of control and test samples were drawn for the control group and test group. The confidence intervals for effect size estimate across bootstrapped resamples are estimated as [bias-corrected and accelerated (BCa) bootstrap confidence intervals](https://blogs.sas.com/content/iml/2017/07/12/bootstrap-bca-interval.html) (Efron, 1987) using a custom implementation in DABEST-Python or boot.ci in dabestr.

#### Mean difference calculation

Mean difference is the bootstrapped mean difference between the control group and the test group:

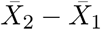

where 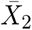 denotes the sample mean of the test group and 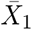 denotes the sample mean of the control group. Mean difference is also calculated for repeated measures, proportion plots, and two-group comparisons in delta-delta and mini-meta designs.

#### Delta-delta mean difference and modified Hedges’ *g* calculations

In the text example for computing delta-delta effects in a 2×2 design over independent variables genotype (“wild type” or “mutant”) and treatment (“placebo” or “drug”), two possible delta-delta effect sizes can be calculated:

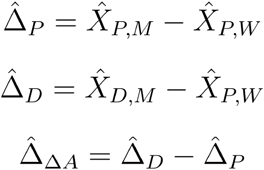

where 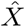denotes the sample mean and Δ denotes the difference of means.

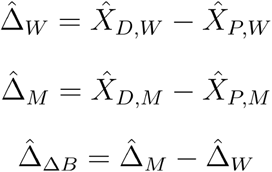

where 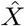 denotes the sample mean and 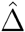 denotes the difference in means.

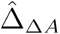 and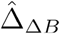 have equal means but distinct variances.

In two-group analysis, standardized effect sizes such as Hedges’ *g* can be computed for comparisons across measurements of different units. For reporting standardized delta-delta effect sizes, DABEST provides a modified Hedges’ *g* calculated from normalization with pooled sample variance.

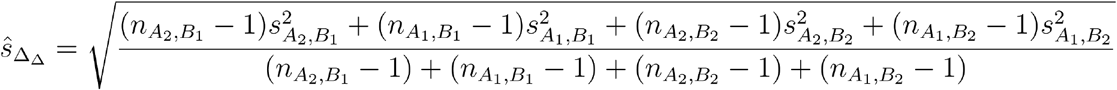

where is the standard deviation and is the sample size.

#### Weighted delta calculation in mini-meta

In a mini-meta experiment, the inverse-variance weighting method is used to calculate a weighted mean difference.

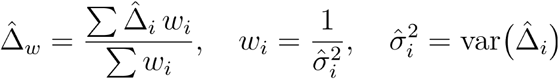

The variance used is calculated as the sample variance of the bootstrapped values of the mean difference.

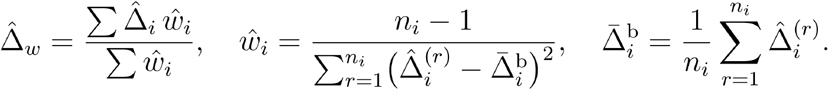

where 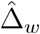 : estimated weighted delta; : weight of mean difference for replicate *i*; 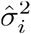: sampling variance of the mean-difference estimator for replicate *i*; 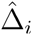: estimated mean difference for replicate *i*; 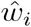: estimated weight of mean difference for replicate *i*; *n_i_*: number of bootstrap replicates used for replicate *i*; 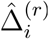: the *r* -th bootstrap estimate of the mean difference for replicate *i*; 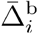: bootstrap mean of the mean differences for replicate *i*

#### Cohen’s *h* calculation for binary data

Cohen’s *h* is an effect size applicable to binary data.

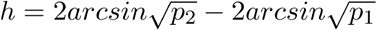

Where denotes the proportion of successes in the test group and denotes that of the control group.

### Visualization

Visualization of the raw data and effect size distributions was achieved with matplotlib and seaborn in DABEST-Python and ggplot2 in dabestr. Except for the Sankey diagram, designs are variations of either the Gardner-Altman plot (Gardner & Altman, 1986; Ho et al., 2019) or the Cumming plot (Cumming, 2011; Ho et al., 2019).

### Sankey Diagrams

An alluvial design of the traditional Sankey plot (Kennedy & Sankey, 1898) was implemented using matplotlib with inspiration from pySankey. A custom data transformation protocol utilized NumPy for numerical computations and pandas for efficient data handling, ensuring precision in managing large datasets. Parameters such as edge thickness and color opacity can be manually adjusted to enhance readability and interpret the dynamic transitions in the data effectively.

### Literature review of proportion statistics usage

A total of 70 articles were sourced to identify the use of Fisher’s exact test in scientific papers. Two types of searches were conducted: the first to break down how Fisher’s exact tests were reported in papers, and the second was to compile a wide range of papers to identify the number of studies that generated proportion data and the visualization associated with it.

The first search (identifying 40 articles) used the keywords “Fisher’s exact test” in PubMed (19,967 publications) and Google Scholar (17,900 publications) from Jan 2000 to May 2024, excluding books and meta-analysis. These articles were further categorized based on the representation of how Fisher’s test results were conveyed, with or without visualizations. The second search (identifying 30 articles) focused on the keyword “Proportion” in PubMed (46,184 publications), also spanning from Jan 2000 to May 2024, and categorized by the presence or absence of error bars/confidence intervals as well as the types of statistical tests that were performed. It was found that about 40% of these papers used Fisher’s exact test, with tables being the predominant form of visualization. Most of the other tests performed were either ANOVA or represented in percentages.

### Software availability

- DABEST is available in both Python and R.
- The Python package, DABEST-python, is hosted on PyPI, and its source code can be found on GitHub at https://github.com/ACCLAB/DABEST-python.
- A tutorial for the Python package can be found at https://acclab.github.io/DABEST-python/tutorials/.
- The R variation, dabestr, is available on CRAN (https://cran.r-project.org/web/packages/dabestr) and similarly has its source code accessible on GitHub (https://github.com/ACCLAB/dabestr).
- A tutorial for the R package is available at https://acclab.github.io/dabestr/.
- Both repositories are released under the Apache-2.0 license.
- In addition, www.estimationstats.com provides an online user interface and performs all the calculations using DABEST-Python.

## Acknowledgements

The authors would like to thank the users of DABEST-python and dabestr for feedback, and Dr Jessica Tamanini and an anonymous biostatistician (Insight Editing London) for critical reviews of the manuscript before submission. We thank Felicia Low for R programming assistance.

